# Impact of JN.1 booster vaccination on neutralisation of SARS-CoV-2 variants KP.3.1.1 and XEC

**DOI:** 10.1101/2024.10.04.616448

**Authors:** Prerna Arora, Christine Happle, Amy Kempf, Inga Nehlmeier, Metodi V. Stankov, Alexandra Dopfer-Jablonka, Georg M. N. Behrens, Stefan Pöhlmann, Markus Hoffmann

## Abstract

The SARS-CoV-2 KP.3.1.1 lineage is currently the dominating lineage on several continents. In parallel, the XEC lineage, a recombinant of KS.1.1 and KP.3.3, is on track of becoming the next dominant lineage in Europe and North America. Here we performed a rapid virological characterisation of the XEC lineage and studied the impact of JN.1 mRNA booster vaccination on KP.3.1.1 and XEC neutralisation.

## Introduction

The SARS-CoV-2 KP.3.1.1 lineage is currently the dominating lineage on several continents. In parallel, the XEC lineage, a recombinant of KS.1.1 and KP.3.3^1^, is rapidly gaining prevalence in Europe and North America, and on track of becoming the next dominant lineage (**Figure 1A**). The SARS-CoV-2 spike (S) protein mediates host cell entry and is the key target of neutralising antibodies. Since the breakpoint of the XEC lineage resides within the S protein gene, resulting in a chimeric S protein harbouring mutations from both KS.1.1 and KP.3.3 (**Figure 1B**), XEC may possess biological traits that enable efficient spread despite high prevalence of KP.3.1.1. For protection against JN.1-derived variants, JN.1-apdated booster vaccines have been developed. However, it is currently unknown whether the vaccine-induced neutralizing antibodies are effective against KP.3.1.1 and XEC.

**Figure 1:**
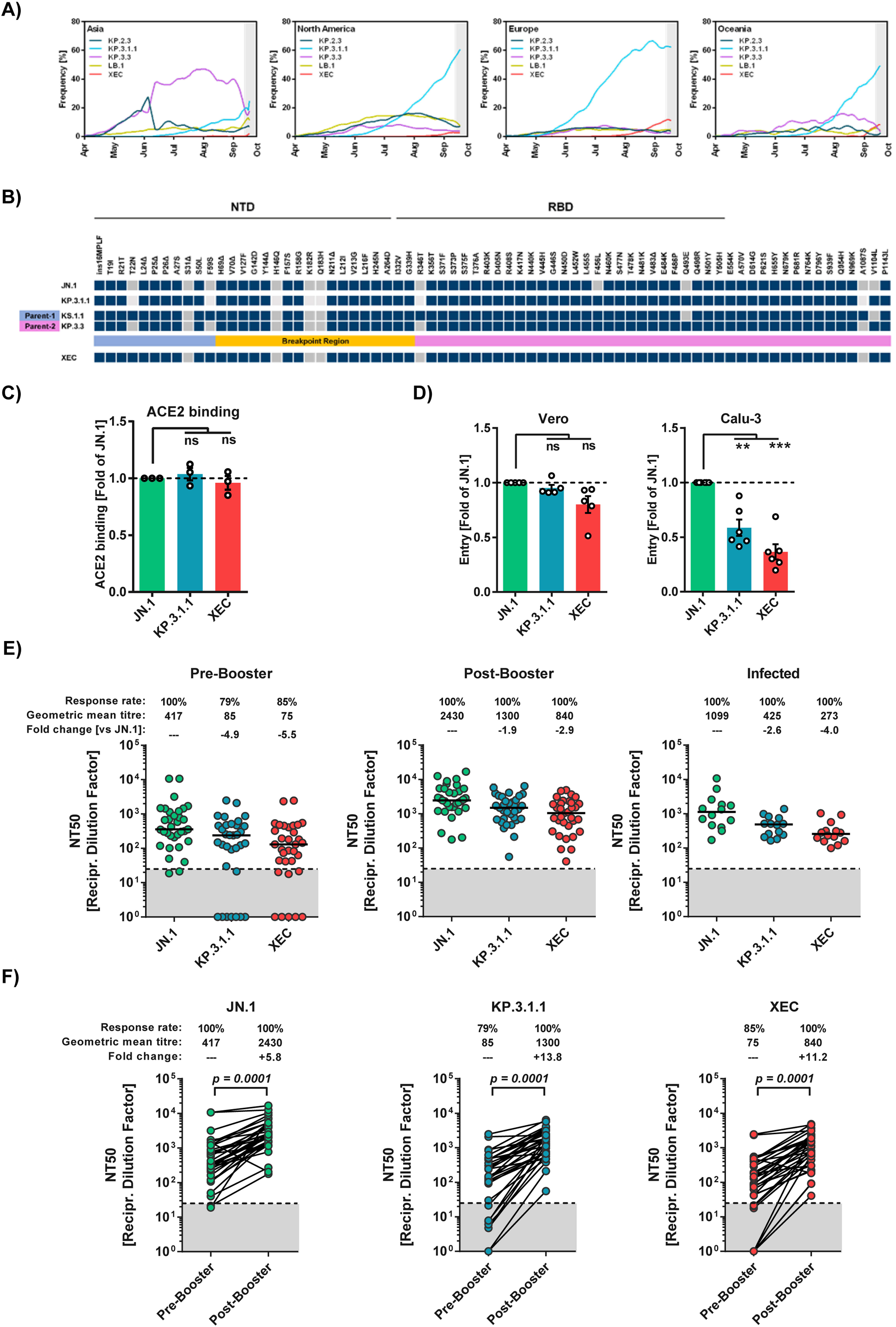
Cell entry and neutralisation escape by SARS-CoV-2 variants KP.3.1.1 and XEC. (A) Relative frequency (seven-day sliding window) of SARS-CoV-2 lineages KP.2.3, KP.3.1.1, KP.3.3, LB.1 and XEC in selected continents (graphs are based on data retrieved from https://cov-spectrum.org/, accessed on 28.09.2024). Grey areas indicate time periods that are prone to underreporting (due to delays between sequencing and uploading of new sequences). (B) Schematic overview of the mutations found in the spike (S) protein of the indicated sublineages (in comparison to the S protein of SARS-CoV-2 Wuhan-Hu-01). Parental lineages (KS.1.1 and KP.3.3) of the recombinant XEC lineage are indicated and the breakpoint region is highlighted in yellow. Abbreviations: NTD = N-terminal domain; RBD = receptor-binding domain. (C) ACE2 binding efficacy of JN.1, KP.3.1.1, and XEC S proteins. 293T cells were transfected with expression plasmids for the indicated S proteins (or no S protein, control) and subsequently analysed by flow cytometry for binding to soluble human ACE2. Presented is the mean of three biological replicates, conducted with a single sample. ACE2 binding was corrected for S protein cell surface expression and normalised using the JN.1 S protein as the reference (=1). Error bars indicate the standard error of the mean (SEM). Statistical significance was assessed by two-tailed Student’s t-test with Welch corrections (p > 0.05, not significant [ns]). Please also see Supplementary Information Figure S2 for more information. (D) Cell entry driven by JN.1, KP.3.1.1, and XEC S proteins. Pseudovirus particles bearing the indicated S proteins were inoculated onto Vero (African green monkey, kidney) and Calu-3 (Human, lung) cells and cell entry was analysed at 16-18 h postinoculation by measuring the activity of virus-encoded firefly luciferase in cell lysates. Presented are the mean data from five (Vero) or six (Calu-3) biological replicates, each performed with four technical replicates. Error bars indicate SEM. Data were normalised against cell entry of JN.1pp (set as 1, dashed line). Statistical significance was assessed by two-tailed Student’s t-test with Welch corrections (p > 0.05, not significant [ns]; p ≤ 0.05, *; p ≤ 0.01, **; p ≤ 0.001, ***). Please also see Supplementary Information Figure S2 for more information. (E) Neutralisation sensitivity of pseudovirus particles bearing JN.1, KP.3.1.1, or XEC S proteins. Two cohorts were analysed: cohort 1 = 33 individuals, sampling was performed before and 21 day after JN.1 booster vaccination; cohort 2 = 14 individuals with recent SARS-CoV-2 infection in the summer of 2024. Pseudovirus particles bearing the indicated S proteins were preincubated with plasma dilutions before being inoculated onto Vero cells. Relative inhibition of pseudovirus entry was calculated using particles incubated in the absence of plasma as control (= 0% inhibition). Individual NT50 (neutralizing titre 50) values were determined for each plasma using a non-linear regression model and geometric mean titres (GMT) were calculated for the respective sample groups. The lowest plasma dilution tested (dashed lines) and the threshold (lower limit of detection, LLOD; grey shaded areas) are indicated. Samples that yielded NT50 values below 12.5 (LLOD) were considered negative and manually assigned a value of 1. Presented are the GMTs (indicated by horizontal black lines and numerical values), response rates and fold changes in neutralisation compared to JN.1pp. Data were derived from a single experiment, performed with four technical replicates. Please also see Supplementary Information Figure S3, S4 and S5 for more information. (F) The donor-matched neutralisation data for cohort 1 before and 21 days after JN.1 booster vaccination are shown (lines connect pre- and post-booster samples from the same plasma donors). Information above the graphs indicate GMTs and the mean fold change in neutralisation between pre- and post-booster samples. Statistical significance was assessed by Wilcoxon matched-pairs signed rank test (p ≤ 0.0001; ****). Please also see Supplementary Information Figure S3, S4 and S5 for more information.

## Results

Here we performed a virological characterisation of the XEC lineage and studied the impact of JN.1-booster vaccination on KP.3.1.1 and XEC neutralisation. We found that the XEC S protein engaged ACE2 with similar efficiency as JN.1 and KP.3.1.1 S proteins (**Figure C**). For analysis of cell entry, we used S protein-bearing pseudovirus particles (pp), which are an established surrogate system to study SARS-CoV-2 cell entry and its neutralisation^2^. Pseudovirus particles bearing JN.1 (JN.1_pp_), KP.3.1.1 (KP.3.1.1_pp_), or XEC S proteins (XEC_pp_) entered Vero kidney cells with comparable efficiency while KP.3.1.1_pp_ and XEC_pp_ exhibited reduced entry into Calu-3 lung cells (**Figure D**). We next investigated whether the new JN.1-adapted mRNA vaccine bretovameran (BioNTech/Pfizer, Mainz, Germany)^3^, which has been demonstrated to booster neutralising activity against JN.1_pp_, KP.2_pp_, KP.2.3_pp_, and LB.1_pp_^4^, would also booster neutralisation of KP.3.1.1_pp_ and XEC_pp_. Our first cohort consisted of 33 healthy individuals and we measured neutralising activity in blood plasma before and 21 days after vaccination. Further, we included another cohort of 14 individuals that had not received the JN.1-booster vaccination but were infected during summer 2024, a period with high prevalence of KP.3.1.1 in Germany. Before JN.1-booster vaccination, neutralisation (given as geometric mean titre, GMT) was highest for JN.1_pp_ (GMT=417), while neutralisation of KP.3.1.1_pp_ (GMT=85) and XEC_pp_ (GMT=75) was reduced by 4.9- and 5.5-fold, respectively (**Figure E**). Three weeks after booster vaccination, neutralisation was increased for all three lineages (5.8-to 13.8-fold, **Figure F**) but neutralisation of KP.3.1.1_pp_ (GMT=1300) and XEC_pp_ (GMT=840) remained lower (1.9- and 2.9-fold, respectively) compared to JN.1_pp_ (GMT=2430) (**Figure E**). For individuals with recent SARS-CoV-2 infection but without JN.1-booster vaccination, neutralisation of JN.1_pp_ (GMT=1099) was 2.6- and 4.0-fold more efficient as compared to KP.3.1.1_pp_ (GMT=425) and XEC_pp_ (GMT=273) (**Figure E**).

## Discussion and Conclusion

Collectively, we found that the XEC S protein retains the ability to efficiently engage ACE2 and drive cell entry, although entry into Calu-3 lung cells was reduced. Both KP.3.1.1_pp_ and XEC_pp_ were generally less well neutralised compared to JN.1_pp,_ indicating elevated immune evasion. Importantly, JN.1-booster vaccination significantly improved neutralisation of all lineages tested and therefore will likely increase protection against hospitalisation and post-COVID sequelae from infection caused by KP.3.1.1 and XEC.

## Acknowledgement

We thank Roberto Cattaneo, Stephan Ludwig, Andrea Maisner, and Gert Zimmer for providing reagents. We gratefully acknowledge the originating laboratories responsible for obtaining the specimens, as well as the submitting laboratories where the genome data were generated and shared via GISAID, on which this research is based. We thank Luise Graichen and Anna-Sophie Moldenhauer for excellent technical assistance. Finally, we thank the participants of the CoCo Study for their support and the entire CoCo study team.

