## Supplemental Material for "Impact of JN.1 booster vaccination on neutralisation of SARS-CoV-2 variants KP.3.1.1 and XEC"

Supplementary figures…………………………………………………..13

### Figure

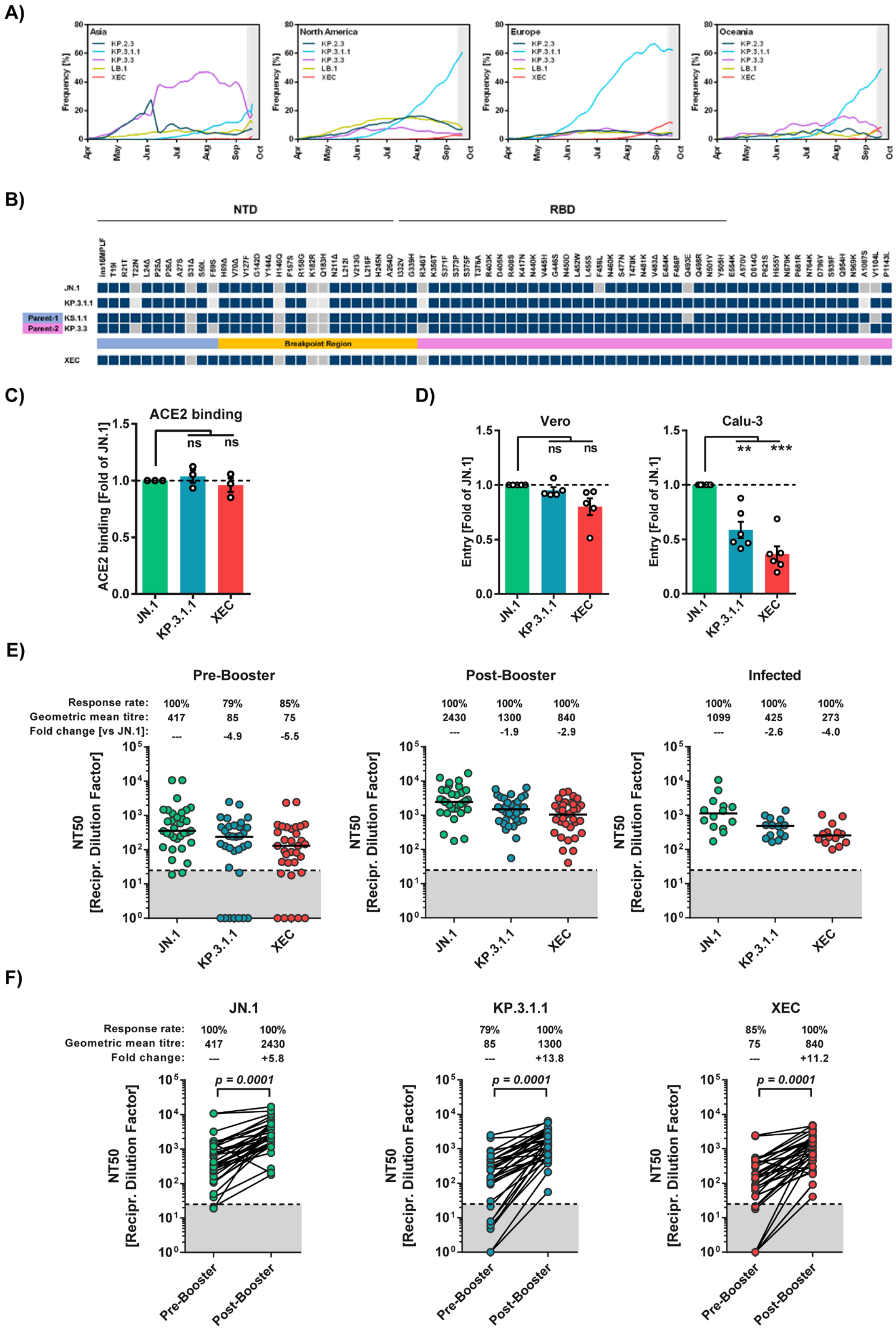

A)
Asia
North America
Europe
Oceania
KP.2.3
KP.3.1.1
KP.3.3
LB.1
XEC
Frequency [%]
Apr
May
Jun
Jul
Aug
Sep
Oct
B)
NTD
RBD
JN.1
KP.3.1.1
Parent-1 KS.1.1
Parent-2 KP.3.3
Breakpoint Region
XEC
C)
ACE2 binding
ACE2 binding [Fold of JN.1]
ns
ns
JN.1
KP.3.1.1
XEC
D)
Vero
Calu-3
Entry [Fold of JN.1]
ns
ns
**
***
E)
Pre-Booster
Post-Booster
Infected
Response rate:
Geometric mean titre:
Fold change [vs JN.1]:
100% 79% 85%
417 85 75
--- -4.9 -5.5
100% 100% 100%
2430 1300 840
--- -1.9 -2.9
100% 100% 100%
1099 425 273
--- -2.6 -4.0
NT50
[Recipr. Dilution Factor]
F)
JN.1
KP.3.1.1
XEC
Response rate:
Geometric mean titre:
Fold change:
100% 100%
417 2430
--- +5.8
79% 100%
85 1300
--- +13.8
85% 100%
75 840
--- +11.2
p = 0.0001
Pre-Booster
Post-Booster

**Figure: Cell entry and neutralisation escape by SARS-CoV-2 variants KP.3.1.1 and XEC.**

(A) Relative frequency (seven-day sliding window) of SARS-CoV-2 lineages KP.2.3, KP.3.1.1, KP.3.3, LB.1 and XEC in selected continents (graphs are based on data retrieved from https://cov-spectrum.org/, accessed on 28.09.2024). Grey areas indicate time periods that are prone to underreporting (due to delays between sequencing and uploading of new sequences).

(B) Schematic overview of the mutations found in the spike (S) protein of the indicated sublineages (in comparison to the S protein of SARS-CoV-2 Wuhan-Hu-01). Parental lineages (KS.1.1 and KP.3.3) of the recombinant XEC lineage are indicated and the breakpoint region is highlighted in yellow. Abbreviations: NTD = N-terminal domain; RBD = receptor-binding domain.

(C) ACE2 binding efficacy of JN.1, KP.3.1.1, and XEC S proteins. 293T cells were transfected with expression plasmids for the indicated S proteins (or no S protein, control) and subsequently analysed by flow cytometry for binding to soluble human ACE2. Presented is the mean of three biological replicates, conducted with a single sample. ACE2 binding was corrected for S protein cell surface expression and normalised using the JN.1 S protein as the reference (=1). Error bars indicate the standard error of the mean (SEM). Statistical significance was assessed by two-tailed Student's t-test with Welch corrections ($p > 0.05$, not significant [ns]). Please also see Supplementary Information Figure S2 for more information.

(D) Cell entry driven by JN.1, KP.3.1.1, and XEC S proteins. Pseudovirus particles bearing the indicated S proteins were inoculated onto Vero (African green monkey, kidney) and Calu-3 (Human, lung) cells and cell entry was analysed at 16-18 h postinoculation by measuring the activity of virus-encoded firefly luciferase in cell lysates. Presented are the mean data from five (Vero) or six (Calu-3) biological replicates, each performed with four technical replicates. Error bars indicate SEM. Data were normalised against cell entry of JN.1pp (set as 1, dashed line).

Statistical significance was assessed by two-tailed Student's t-test with Welch corrections ($p > 0.05$, not significant [ns]; $p \leq 0.05$, *; $p \leq 0.01$, **; $p \leq 0.001$, ***). Please also see Supplementary Information Figure S2 for more information.

(E) Neutralisation sensitivity of pseudovirus particles bearing JN.1, KP.3.1.1, or XEC S proteins. Two cohorts were analysed: cohort 1 = 33 individuals, sampling was performed before and 21 day after JN.1 booster vaccination; cohort 2 = 14 individuals with recent SARS-CoV-2 infection in the summer of 2024. Pseudovirus particles bearing the indicated S proteins were preincubated with plasma dilutions before being inoculated onto Vero cells. Relative inhibition of pseudovirus entry was calculated using particles incubated in the absence of plasma as control (= 0% inhibition). Individual NT50 (neutralizing titre 50) values were determined for each plasma using a non-linear regression model and geometric mean titres (GMT) were calculated for the respective sample groups. The lowest plasma dilution tested (dashed lines) and the threshold (lower limit of detection, LLOD; grey shaded areas) are indicated. Samples that yielded NT50 values below 12·5 (LLOD) were considered negative and manually assigned a value of 1. Presented are the GMTs (indicated by horizontal black lines and numerical values), response rates and fold changes in neutralisation compared to JN.1pp. Data were derived from a single experiment, performed with four technical replicates. Please also see Supplementary Information Figure S3, S4 and S5 for more information.

(F) The donor-matched neutralisation data for cohort 1 before and 21 days after JN.1 booster vaccination are shown (lines connect pre- and post-booster samples from the same plasma donors). Information above the graphs indicate GMTs and the mean fold change in neutralisation between pre- and post-booster samples. Statistical significance was assessed by Wilcoxon matched-pairs signed rank test ($p \leq 0.0001$; ****). Please also see Supplementary Information Figure S3, S4 and S5 for more information.

**Table S1: Demographics, infection and vaccination history**

| Variable | Vaccinees (cohort 1) | Breakthrough infections (cohort 2) |
|---|---|---|
| Study participants (n=) | 33 | 14 |
| Age, Median [IQR] (years) | 45 [24] | 45 [28] |
| Sex, male (%) | 45·5 | 21 |
| Median time post last vaccination [IQR] (months) | 11 [6·8] | 28 [22] |
| Median number of prior vaccinations [IQR] | 4 [1] | 4 [2] |
| Prior SARS-CoV-2 omicron vaccination (%) | 87·9 | 69·2 |
| Prior SARS-CoV-2 infection (%) | 93·3 | 100 |
| Prior SARS-CoV-2 omicron infection (%) | 90·3 | 100 |
| Prior SARS-CoV-2 omicron antigen contact (%) | 100 | 100 |

Abbreviation: IQR, interquartile range

#### Methods

##### Cell culture

All cell lines were cultured in a humidified environment with 5% $CO_2$ at 37 °C. 293T (human, female, kidney; ACC-635, DSMZ; RRID: CVCL_0063) and Vero cells (African green monkey kidney, female, kidney; CRL-1586, ATCC; RRID: CVCL_0574, kindly provided by Andrea Maisner) were grown in Dulbecco's Modified Eagle Medium (DMEM, PAN-Biotech) supplemented with 10% fetal bovine serum (FBS, Biochrom), 100 U/ml of penicillin, and 0.1 mg/ml of streptomycin (P/S, PAN-Biotech). Calu-3 (human, male, lung; HTB-55, ATCC; RRID: CVCL_0609, kindly provided by Stephan Ludwig) were cultured in Dulbecco's Modified Eagle Medium F12 (DMEM/F-12, GIBCO, supplemented with 10% FBS, 1% P/S, non-essential amino acid solution (NEAA, PAN-Biotech, 1:100 v/v dilution), and 1 mM sodium pyruvate (PAN-Biotech). Cell lines were validated using STR analysis, amplification and sequencing of a cytochrome c oxidase gene fragment, microscopic investigation, and/or growth characteristics. Moreover, all cell lines underwent routine testing for mycoplasma contamination. 293T cells were transfected using the calcium phosphate method.

##### Expression plasmids and sequence analysis

The expression plasmids pCAGGS-DsRed, pCG1-sol-ACE2-Fc, pCG1-SARS-CoV-2 JN.1 SΔ18 (codon-optimised with a C-terminal truncation of 18 amino acids, GISAID Accession ID: EPI_ISL_18530042) have previously been described elsewhere [1,2]. We employed Gibson assembly to generate expression plasmids for SARS-CoV-2 KP.3.1.1 S Δ18 (codon-optimised with a C-terminal truncation of 18 amino acids, GISAID Accession ID: EPI_ISL_EPI_ISL_19455032) and SARS-CoV-2 XEC S Δ18 (codon-optimised with a C-terminal truncation of 18 amino acids, GISAID Accession ID: EPI_ISL_19454087) using one overlapping DNA string (Thermo Fisher Scientific), BamHI/XbaI-digested pCG1 plasmid and GeneArt™ Gibson Assembly HiFi Master Mix (Thermo Fisher Scientific). The reactions were prepared according to the manufacturer's instructions. The pCG1 expression plasmid was generously provided by Roberto Cattaneo from the Mayo Clinic College of Medicine, Rochester, MN, USA. All PCR-amplified sequences were verified using a commercial sequencing service (Microsynth SeqLab). Data regarding SARS-CoV-2 lineages and spike protein sequences were retrieved from the GISAID (Global Initiative on Sharing All Influenza Data) and CoV-Spectrum (https://cov-spectrum.org/) databases, accessed on 28.09.2024.

**Pseudovirus particle production**

Pseudovirus particles containing SARS-CoV-2 S proteins were generated as previously described [3]. In brief, 293T cells transfected to express the respective S protein or DsRed (control) were infected with VSV-G-trans-complemented VSV*G (FLuc) generously provided by Gert Zimmer [4]. After one hour of incubation, the supernatant was aspirated, and cells were washed once with phosphate-buffered saline (PBS). The cells were then incubated for an additional 16–18 hours in DMEM medium containing an anti-VSV-G antibody (culture supernatant from I1-hybridoma cells; ATCC no CRL-2700) expected for cells producing pseudovirus particles harbouring VSV-G. Culture supernatants containing pseudovirus particles were harvested, cleared from cellular debris by centrifugation (4,000 × g, 10 min), and directly used for cell entry and neutralisation experiments or stored at -80 °C for further use.

**Transduction of target cells**

Target cells were seeded in 96-well plates and equivalent amounts of pseudovirus particles were used to transduce the cells. The transduction efficiency was assessed by measuring luciferase activity in cell lysates at 16–18 hours post-transduction. The cells were lysed using PBS with 0.5% Tergitol (Carl Roth) for 30 minutes at room temperature. Subsequently, a luciferase substrate (Beetle-Juice, PJK) was added to the cell lysates in white 96-well plates, and luminescence was measured using a Hidex Sense plate luminometer (Hidex).

**Analysis of spike protein cell surface expression and ACE2 binding efficiency**

Cells were either transfected with plasmids encoding S proteins or an empty control plasmid, culture supernatants were removed, and cells were resuspended in FACS buffer (PBS with 1% BSA) and split into two tubes for the detection of S protein cell surface expression and ACE2 binding. (i) For S protein cell surface expression, cells were incubated with anti-SARS-CoV-2 S2 subunit antibody (mouse, 1:100 in FACS buffer; GTX632604, Biozol) for 1 h at 4 °C. Thereafter, the cells were pelleted by centrifugation and washed with FACS buffer before they were incubated for 1 h at 4 °C with Alexa Fluor-488-conjugated anti-mouse antibody (1:200 in PBS-B; A-10667, Thermo Fisher Scientific). (ii) For ACE2 binding, cells were incubated for with soluble human ACE2-Fc (concentrated supernatant of 293T cells transfected with pCG1-sol-ACE2-Fc; 1:20 in FACS buffer) for 1 h at 4 °C. Thereafter, the cells were pelleted by centrifugation and washed with FACS-buffer before they were incubated for 1 h at 4 °C with Alexa Fluor-488-conjugated anti-human antibody (1:200 in FACS buffer; A-11013, Thermo Fisher Scientific). For both (i) and (ii), the cells were pelleted by centrifugation, washed twice with FACS-buffer and resuspended in FACS buffer before S protein cell surface expression and ACE2 binding was analysed using an ID7000 Spectral Cell Analyzer (Sony Biotechnology) and ID7000 software. Finally, for each S protein, ACE2 binding was normalised to S protein surface expression. All centrifugation steps described above were carried out at 600× g, 5 min, room temperature.

**Plasma collection and neutralisation assay**

For this analysis, we considered individuals without underlying conditions or therapy with potentially immunomodulatory effects from the COVID-19 Contact (CoCo) Study, which comprises a general population of health-care professionals at Hannover Medical School. We report about n=33 individuals, constituting cohort 1, that were vaccinated with Comirnaty® omicron JN.1/bretovameran, for which baseline and 21-day follow-up data was available and which had no evidence for a SARS-CoV-2 infection between vaccination and day 21 (no self-reported positive quick test/PCR or new positive anti-nucleocapsid protein (NCP) IgG). In addition, we included as cohort 2 n=14 participants with recent COVID-19 in July or August 2024 confirmed by quick test for comparison (all anti-NCP IgG positive). According to the German wastewater-based surveillance on SARS-CoV-2, KP.3.1.1, LB.1, KP.3, KP.2 and JN.1 were among the most detected variants [5]. Anti SARS-CoV-2 IgG titres were measured using the anti-SARS-CoV-2-QuantiVac and anti-SARS-CoV-2 NCP IgG ELISA(EUROIMMUN, Lübeck, Germany). More information on demographics, COVID-19 vaccine and infection history of participants is provided in Supplementary Table S1. None of the participants with a history of

SARS-CoV-2 infection experienced severe disease. Plasma samples were heat-inactivated at 56 °C for 30 minutes prior to experimentation.

Neutralisation assays were carried out as described before [6]. Briefly, particles containing the respective S proteins were mixed with dilutions of blood plasma ranging from 1:25 to 1:25,600, and incubated for 30 minutes at 37 °C. Subsequently, these mixtures were added to Vero cells. After an incubation period of 16 to 18 hours, the neutralisation efficiency was assessed. For this, cell entry of pseudovirus particles was measured relative to samples without plasma, which was set as 0% inhibition. The concentration of plasma dilution resulting in half-maximal inhibition (neutralizing titre 50, $NT_{50}$ for plasma) was determined using a non-linear regression model. The thresholds for neutralisation positivity were defined as $NT_{50} \geq 12.5$ (50% of the lowest plasma dilution tested).

**Statistical analysis**

Microsoft Excel (part of Microsoft Office Professional Plus, version 2016, Microsoft Corporation) and GraphPad Prism version 6.07 were used to analyse the data (GraphPad Software). The two-tailed Students t-test (cell entry mediated by S protein, ACE2 binding) or Wilcoxon signed-rank test (plasma neutralisation) were used to determine statistical significance. Only p-values of 0.05 or less were considered statistically significant ($p > 0.05$, not significant [ns], $p \leq 0.05$, *, $p \leq 0.01$, **, $p \leq 0.001$, ***).

**Ethics committee approval**

The collection and analysis of plasma samples was conducted as part of the COVID-19 Contact (CoCo) Study (DRKS00021152), approved by Hannover Medical School's Internal Review Board (no. 8973_BO_K_2020). This prospective observational study monitors IgG and immune responses in healthcare professionals and individuals with potential SARS-CoV-2 contact at Hannover Medical School. Participants provided informed consent and received no compensation.

#### Limitations of the study

This study has limitations. First, pseudovirus particles were employed as a surrogate model to investigate neutralisation of SARS-CoV-2 variants. While these pseudovirus particles adequately mirror neutralisation of authentic SARS-CoV-2 by antibodies [7], formally our findings require validation with authentic virus. Second, the data presented here offer only an initial understanding of the immune response elicited by the updated JN.1 vaccine. Thus, long-term studies are needed to evaluate the durability and evolving nature of this immune response. Third, the majority of participants in cohort 1 had on or more prior SARS-CoV-2 infections and/or multiple vaccinations, which could have affected humoral immune responses. Fourth, XEC harbours mutations outside the S gene that may impact viral fitness but could not be analysed using our pseudovirus particle system.

##### Acknowledgements

We thank Anna-Sophie Moldenhauer, Andrea Stölting, Simon F. Ritter, Louis Kuhnke, Eva Stöppelmann and Luis Manthey for technical help. We would want to express our gratitude to the originating laboratories who were in charge of collecting the samples as well as the submitting laboratories that produced and transmitted the genome data via GISAID that served as the foundation for our study.

### Supplementary figures

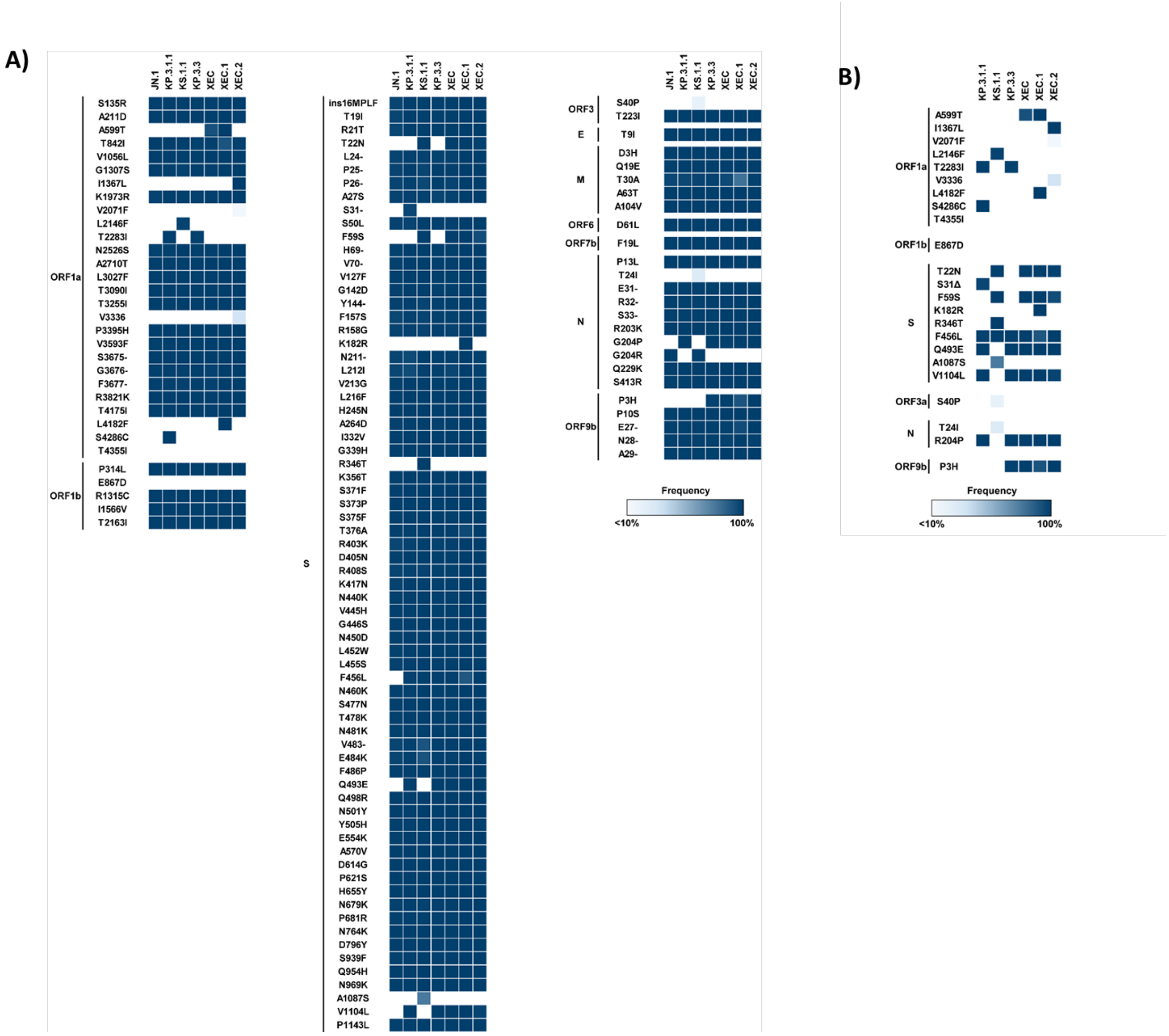

**Supplementary figure 1: Whole genome missense mutations in comparison to Wuhan-01**

Whole genome missense mutations in JN.1 and JN.1-derived variants in comparison to Wuhan-01 (A) or JN.1 (B). The underlying data were retrieved from https://cov-spectrum.org/ (accessed on 28.09.2024).

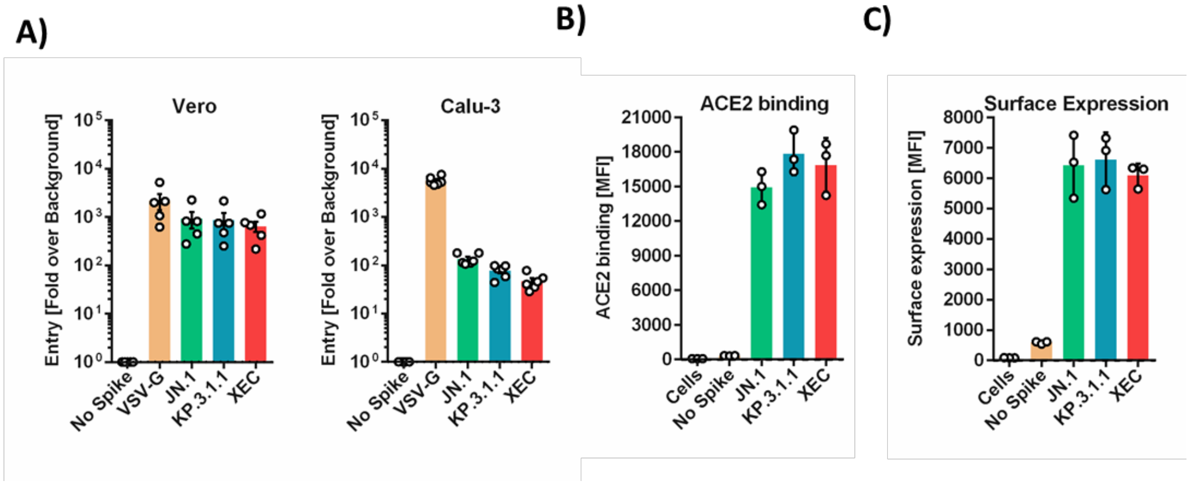

**Supplementary figure 2: Cell tropism, ACE2 binding and surface expression.**

(A) The pseudovirus entry data were normalised against the assay background (luciferase activity obtained for particles bearing no viral surface protein, set as 1). Further, data for particles bearing vesicular stomatitis virus glycoprotein (VSV-G) are included.

(B) 293T cells expressing the indicated S proteins following transfection were incubated with soluble human ACE2-Fc and Alexa Fluor-488-conjugated anti-human antibody, before ACE2 binding was analysed by flow cytometry. Presented are the mean fluorescence intensity (MFI) data from three biological replicates. Error bars indicate standard deviation (SD).

(C) 293T cells expressing the indicated S proteins following transfection were incubated with anti-SARS-CoV-2 S protein S2 subunit and Alexa Fluor-488-conjugated anti-mouse secondary antibodies, before S protein surface expression was analysed by flow cytometry. Presented are the mean MFI data from three biological replicates. Error bars indicate SD..

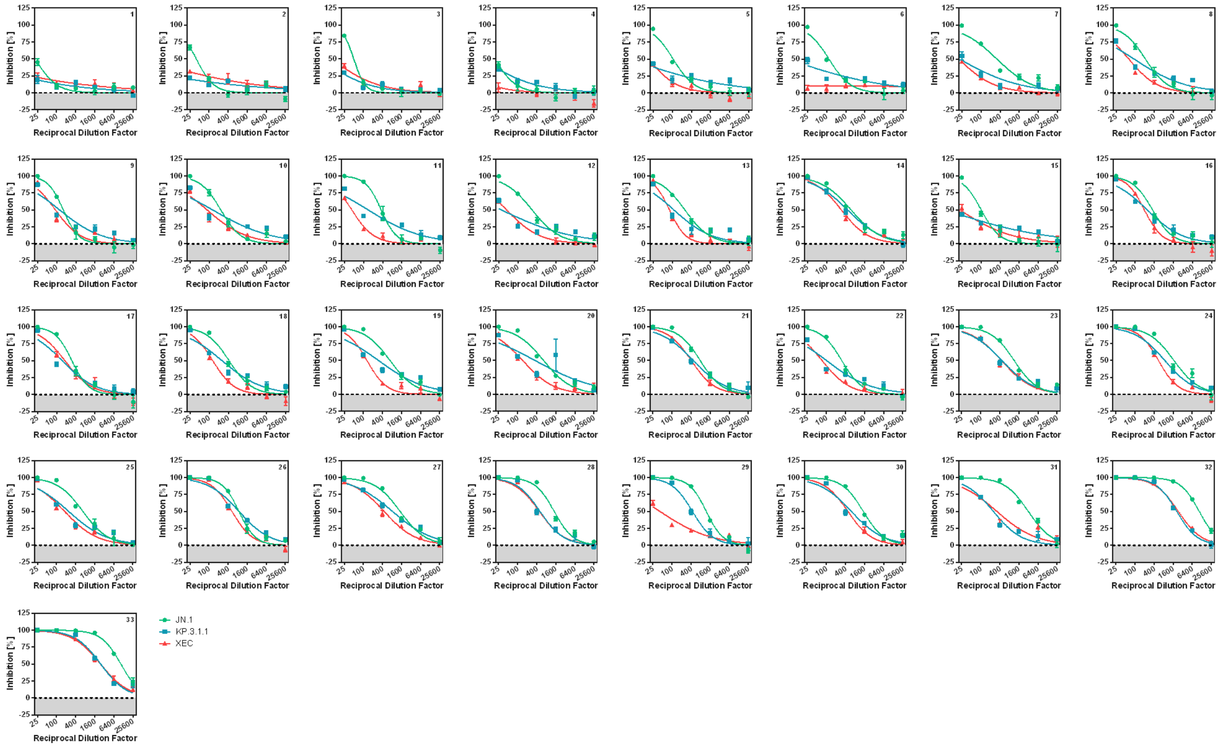

**Supplementary figure 3: Individual neutralisation data for cohort 1 before JN.1 booster vaccination.**

Individual data on JN.$1_{pp}$, KP.3.1.$1_{pp}$ and $XEC_{pp}$ neutralisation by antibodies present in in the blood plasma of individuals before JN.1 booster vaccination. Pseudovirus particles bearing the indicated S proteins were preincubated with serial dilutions of plasma before being inoculated onto Vero cells. Pseudovirus particles incubated in the absence of plasma served as control. Presented are the mean data from a single experiment, performed with four technical replicates. Error bars indicate the standard deviation. S protein-driven cell entry was analysed and normalised to samples without plasma (= 0% inhibition).

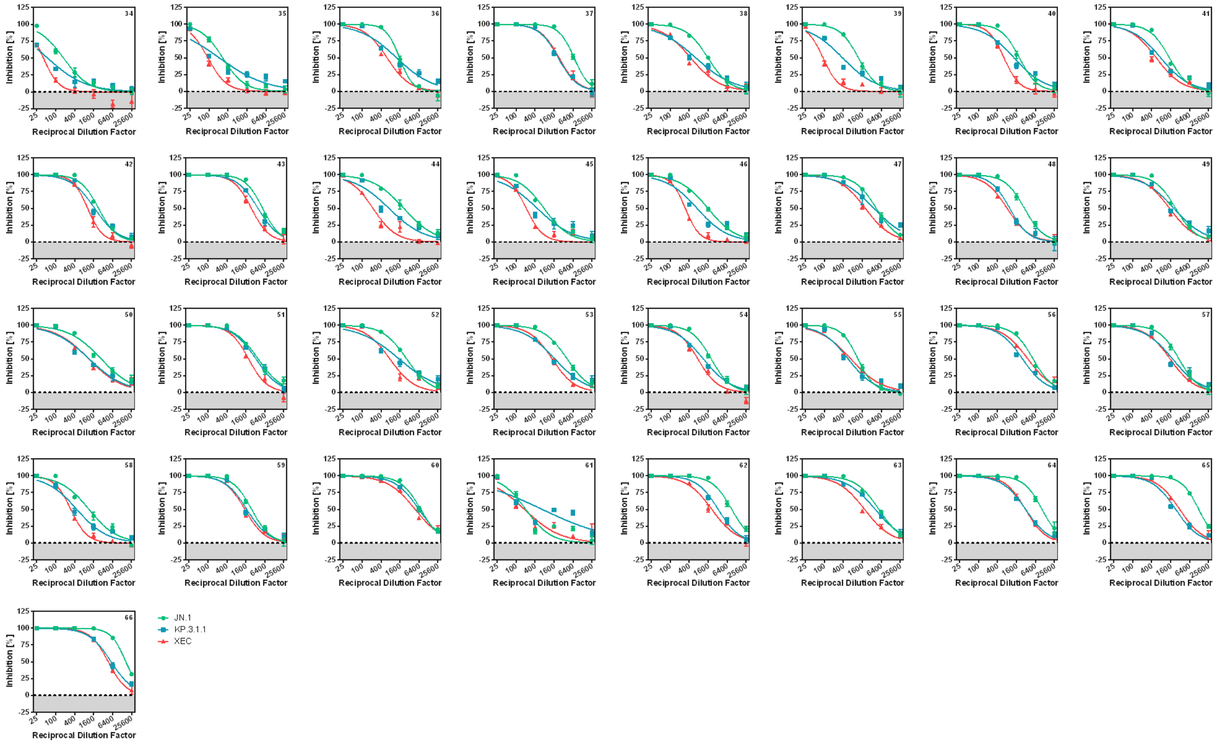

**Supplementary figure 4: Individual neutralisation data for cohort 1 at 21 days post JN.1 booster vaccination.**

Individual data on JN.1$_{pp}$, KP.3.1.1$_{pp}$ and XEC$_{pp}$ neutralisation by antibodies present in the blood plasma of individuals at 21 days post JN.1 booster vaccination. Pseudovirus particles bearing the indicated S proteins were preincubated with serial dilutions of plasma before being inoculated onto Vero cells. Pseudovirus particles incubated in the absence of plasma served as control. Presented are the mean data from a single experiment, performed with four technical replicates. Error bars indicate the standard deviation. S protein-driven cell entry was analysed and normalised to samples without plasma (= 0% inhibition).

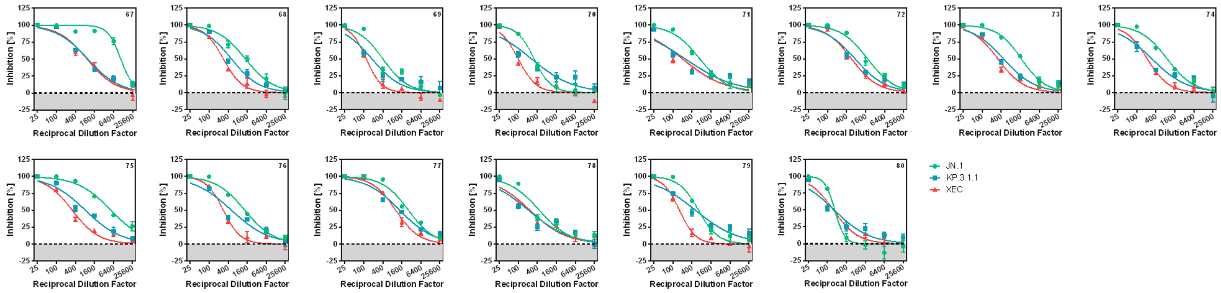

**Supplementary figure 5: Individual neutralisation data for cohort 2.**

Individual data for JN.$1_{pp}$, KP.3.1.$1_{pp}$ and $XEC_{pp}$ neutralisation by antibodies present in in the blood plasma of individuals with SARS-CoV-2 infection in the summer of 2024 in Germany. Pseudovirus particles bearing the indicated S proteins were preincubated with serial dilutions of plasma before being inoculated onto Vero cells. Pseudovirus particles incubated in the absence of plasma served as control. Presented are the mean data from a single experiment, performed with four technical replicates. Error bars indicate the standard deviation. S protein-driven cell entry was analysed and normalised to samples without plasma (= 0% inhibition).
